# Towards semantic interoperability: finding and repairing hidden contradictions in biomedical ontologies

**DOI:** 10.1101/2020.05.16.099309

**Authors:** Luke T Slater, Georgios V Gkoutos, Robert Hoehndorf

**Affiliations:** College of Medical and Dental Sciences, Institute of Cancer and Genomic Sciences, University of Birmingham; Institute of Translational Medicine, University Hospitals Birmingham, NHS Foundation Trust; NIHR Experimental Cancer Medicine Centre; NIHR Surgical Reconstruction and Microbiology Research Centre; NIHR Biomedical Research Centre; MRC Health Data Research UK (HDR UK) Midlands; Computer, Electrical and Mathematical Sciences & Engineering Division, Computational Bioscience Research Center, King Abdullah University of Science and Technology

## Abstract

**Background:** Ontologies are widely used throughout the biomedical domain. These ontologies formally represent the classes and relations assumed to exist within a domain. As scientific domains are deeply interlinked, so too are their representations. While individual ontologies can be tested for consistency and coherency using automated reasoning methods, systematically combining ontologies of multiple domains together may reveal previously hidden contradictions.

**Results:** We developed a method that tests for hidden unsatisfiabilities in an ontology that arise when combined with other ontologies. For this purpose, we combine sets of ontologies and use automated reasoning to determine whether unsatisfiable classes are present. We test the mutual consistency of the OBO Foundry and the OBO ontologies and find that the combined OBO Foundry gives rise to at least 636 unsatisfiable classes, while the OBO ontologies give rise to more than 300,000 unsatisfiable classes.

We design and implement a novel algorithm that can determine justifications for contradictions across extremely large and complicated ontologies, and use these justifications to semi-automatically repair ontologies by identifying the minimal set of axioms that, when removed, result in a consistent and coherent set of ontologies. We applied our algorithm to each combination of OBO ontologies that resulted in unsatisfiable classes.

**Conclusions:** We identified a large set of hidden unsatisfiability across a broad range of biomedical ontologies, and we find that this large set of unsatisfiable classes is the result of a relatively small amount of axiomatic disagreements. Our results show that hidden unsatisfiability is a serious problem in ontology interoperability; however, our results also provide a way towards more consistent ontologies by addressing the issues we identified.

## Introduction

Ontologies are used to describe and organise domain knowledge in the biomedical sciences. Ontologies use classes to characterise the kinds of things that exist within a domain as well as axioms that provide constraints for these classes and conditions that must be satisfied within the domain. Most ontologies in biology are domain-specific and focus on a single domain. Creating ontologies that reference and extend other biomedical ontologies is common practice, as it promotes a unified understanding of the biomedical domain by defining terms and groups of terms in the context of their relationships with classes from related domains, and in the common context of higher level domains. Reusing the formalised knowledge from other domain ontologies also enables the reuse of expertise from ontology developers in other domains.

The majority of biomedical ontologies are now being developed in the Web Ontology Language (OWL) [1], a formal model-theoretic language based on description logics [2]. OWL ontologies enable the use of automated reasoners, which in turn enable the deductive inference of knowledge implied by the explicit assertions made in the ontologies. Furthermore, these inferences can be examined to determine whether an ontology’s classes are satisfiable, and whether an ontology is consistent. A class is satisfiable if it can have an instance, and is unsatisfiable if it contains a contradiction such that an instance of the class would force any model of the ontology to contain a logical contradiction; an ontology is inconsistent if it contains at least one instance of a logical contradiction. Unsatisfiable classes and inconsistencies arise most frequently by violation of a disjointness axiom. For example, if an ontology contains an axiom asserting that a disease and a phenotype are disjoint, then any class that is a subclass of both disease and phenotype is unsatisfiable. An ontology which contains any instances of an unsatisfiable class is inconsistent, while an ontology which contains any unsatisfiable classes is termed incoherent.

Automated reasoners can also be used to generate explanations for an unsatisfiability. An explanation is a small set of axioms which are sufficient to reproduce the contradiction. An explanation can be used to diagnose the cause of the class becoming unsatisfiable.

The Open Biomedical Ontologies (OBO) Foundry is a collection of ontologies that use a shared set of design principles, and encourages re-use of terms amongst them [3]. The ontologies are built using the framework provided by common upper-level ontology, the Basic Formal Ontology (BFO) [4], and include many large and widely used domain ontologies describing areas such as chemical entities [5], phenotypes [6], and model organisms [7]. Using standard upper-level ontologies is intended to support consistency between multiple ontologies and knowledge integration across domains [8].

From a technical perspective, OWL caters for the inclusion (i.e., import) of complete ontologies so that they can be reused and built upon. Importing an ontology amounts to including all the entities and axioms of another ontology in the importing ontology. While this is a provision of simple modularity, it enables re-use of classes and axioms across ontologies, and it enables automated reasoners to detect joint consistency.

However, full import of an ontology is not always sensible or feasible. Even when an ontology makes heavy use of the classes and axioms in another ontology, only a subset of the classes are likely to be relevant within another ontology.

For example, the Hypertension Ontology (HTN) [9] expands upon the hypertension classes in the Human Phenotype Ontology (HP) [6] and the Disease Ontology (DO) [10], but is not concerned with any terms in those ontologies besides those directly related to hypertension. To include all of the classes in HP and DO in HTN is vulnerable to potential issues resulting from the inclusion of irrelevant classes. Loading the ontology would take longer, in particular when imported ontologies are retrieved over the internet. Editing an ontology may become challenging when many classes from other ontologies are included on account of the large amount of additional classes that must be loaded, classified, and possibly visualised. Overall, an ontology importing a large number of other ontologies becomes more difficult to use with the relevant classes being hidden within the hierarchy of the imported ontologies.

In response to these technical challenges, the research community has investigated different models for ontology modularisation. Particularly, work has investigated locality-based module extraction [11], which can be used to improve reasoner-based query performance and support large-scale ontology development and re-use [12].

The MIREOT (Minimum Information to Reference an External Ontology Term) guidelines were originally developed to support inclusion of classes from non-OBO Foundry ontologies without needing to align to their axiomatisation, and has become a standard for term re-use and inclusion throughout the biomedical ontology community [13].

MIREOT relaxes the import of other ontologies through including all axioms and instead focuses on the reuse of individual classes from other ontologies. Particularly, the MIREOT guidelines stipulate that three pieces of information are necessary to “reference” an external ontology class:

**Source ontology** The Internationalised Resource Identifier (IRI) of the ontology which contains the class being included.
**Source class** The IRI of the class to import, as defined in the external ontology.
**Direct Superclass** The IRI of the direct superclass of the imported class in the referencing ontology.

Utilizing these three pieces of information, an external ontology class can be referenced. By including MIREOT definitions for each relevant external class, a module is formed within the imported ontology without fully importing any external ontologies. While this method allows ontologies to reuse classes in a scalable and efficient manner, the inclusion of external classes without the context of the external ontology’s axioms means that contradictions may arise that cannot be detected using an automated reasoner that evaluates only the importing ontology. This may lead ontology developers to build upon another class in a way that contradicts its original definition. Furthermore, subsequent versions of the source ontology may re-axiomatise a subject class in a way which renders its use in the importing ontology incompatible with it.

Our prior analysis of the Experimental Factor Ontology (EFO) [14] showed that the use of MIREOT has the potential to cause inconsistency and unsatisfiabilities across the set of ontologies the EFO references [15]. While our previous work revealed problems with EFO, the extent and exact characterisation of this problem throughout the entire biomedical ontology ecosystem has not yet been explored. It is also unknown whether there are common roots to widespread unsatisfiabilities. More importantly, while identifying unsatisfiable classes and inconsistencies is important, it would be much more useful to resolve them, ideally automatically or semi-automatically. It is not clear whether the unsatisfiabilities can be automatically repaired.

We explore interoperability and hidden unsatisfiability throughout the OBO Foundry ontologies. To do this extend the unMIREOT tool described by our previous work, and generalise it to reveal hidden contradictions in any combination of OWL ontologies [15]. This analysis reveals many cases of incoherency and inconsistency throughout the ontology ecosystem.

Based on the information revealed by our analysis, we present a novel algorithm that generates explanations for unsatisfiability, and uses these explanations to systematically identify a small list of axioms that can be removed from an ontology to repair all cases of unsatisfiability and generate a novel ontology that is both consistent and coherent. The list is formed by automatically evaluating explanations for unsatisfiable classes. We then use the algorithm to report on sources of the contradictions we found throughout the OBO ontologies, and the axioms that are most frequently involved.

Our method and tools allows detection of unsatisfiable classes and the systematic, semi-automatic repair of ontologies. Applying our approach will lead to higher quality ontologies maintaining consistency in the rapidly evolving web of knowledge that spans biology and biomedicine. All our results and software are freely available at https://github.com/bio-ontology-research-group/UNMIREOT.

## Materials and Methods

### Ontologies and ontology versions

All non-deprecated and obtainable OBO ontologies were downloaded using the permanent download links given by the OBO Foundry database at http://obofoundry.org/registry/ontologies.yml. A total of 132 ontologies were obtained on 28/03/2018.

Our experiments concern two sets of ontologies described by this database. First, the OBO Foundry ontologies, which are judged as satisfying the OBO Foundry principles, and are therefore tightly integrated and also widely used across many domains. The second is the wider set of ontologies included in the OBO database. In the rest of this paper, we will refer to the core ontologies as the OBO Foundry ontologies, while the wider set of ontologies will be referred to as the OBO ontologies.

### Implementation and experimental Setup

For all experiments, we use the OWLAPI 5.1.4 [16]. To classify the ontologies and to retrieve unsatisfiability explanations, we use the Elk reasoner version 0.5.0-SNAPSHOT [17].

Elk supports the OWL 2 EL profile, a fragment of OWL that supports tractable (i.e., polynomial-time) reasoning, but lacks support for many logic operators. In particular, OWL 2 EL does not support the use of negation in class descriptions or use of the universal quantifier. The only type of axiom in OWL 2 EL that could result in an explicit contradiction is the disjointness axiom. We also used Protégé to examine some of the combined ontologies for particular cases of unsatisfiability [18].

## Results

### Combining ontologies and detecting inconsistencies

We combined all of the OBO Foundry ontologies into one meta-ontology, by copying all the axioms from each source ontology into the a new ontology. Figure 1 summarises the ontologies in this set. We did not include the ontologies referenced in the imports closures of the OBO Foundry ontologies, since in all cases these ontologies were included in the larger set of OBO ontologies, and therefore their combined consistency would be evaluated later. Subsequently, we evaluated the combined ontology for unsatisfiability and its causes.

**Figure 1.**
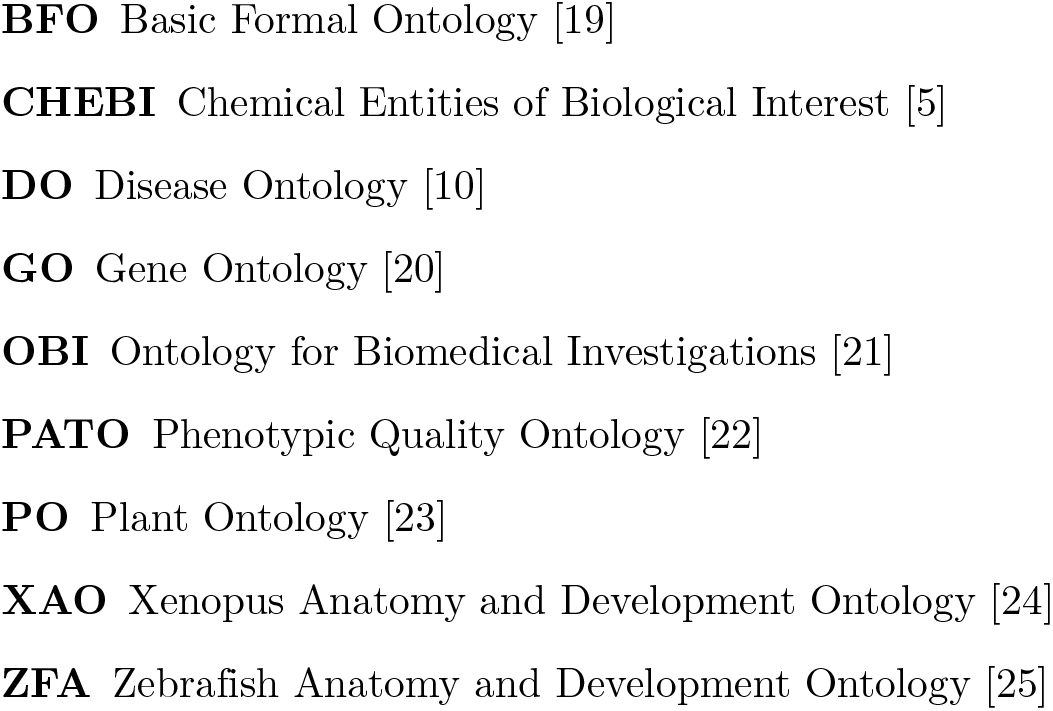
Ontologies included in the OBO Foundry.

The 9 OBO Foundry ontologies combined consist of 402,868 logical axioms and 207,105 named classes. The use of an automated reasoner on the combined OBO Foundry meta-ontology determined that 636 of these classes are unsatisfiable. Table 1 shows the number of unsatisfiable classes and the ontology to which they belong. The origin ontology of the classes was determined using the class IRI prefix.

**Table 1.** Unsatisfiable class counts in OBO Foundry

| Ontology | Unsatisfiable Class Count |
| --- | --- |
| CHEBI | 37 |
| GO | 565 |
| OBI | 34 |

While each of these classes is unsatisfiable due to a different set of axioms, there may be a small set of axioms that are shared by several cases of unsatisfiability. We developed an algorithm to identify a small set of axioms that are sufficient to explain all unsatisfiable classes in an ontology; if this set of axioms is removed from an ontology, all cases of unsatisfiability are resolved. We apply this algorithm to the combined OBO Foundry ontologies in order to derive a coherent version, removing two axioms. The algorithm, and the axioms it removes, are described in detail in the *Efficient ranking and repairing of axioms* section.

We combine this coherent version of the OBO Foundry meta-ontology iteratively with each of the OBO ontologies, classifying the resulting merged ontology, using an automated reasoner to determine if there are any unsatisfiable classes; if we identify unsatisfiable classes we count their number. Out of all 131 loadable ontologies that we use in this experiment, we revealed unsatisfiable classes in 50 ontologies. The 10 OBO ontologies with the most unsatisfiable classes are listed in Table 2. The total number of unsatisfiable classes across all OBO ontologies is 866,494 and the total number of unique unsatisfiable classes is 312,398. Of these, 8,893 are obsolete classes, which are intentionally unsatisfiable (and thus not considered an error). In addition, the Ontology of Vaccine Adverse Events (OVAE) [26], Food Ontology (FOODON) [27], Plant Trait Ontology (TO) [28], Gazetteer (GAZ) [29], Porifera (PORO) [30], Plant Experimental Conditions Ontology (PECO) [28], Oral Health and Disease Ontology (OHD) [31], and Statistics Ontology (STATO) [32] became inconsistent.

**Table 2.** The ten ontologies with the most unsatisfiable classes in the OBO ontologies, when combined with a repaired version of the merged OBO Foundry ontology.

| Ontology Name | Unsatisfiable Class Count |
| --- | --- |
| Unified Phenotype Ontology (UPHENO) [33] | 106,126 |
| Monarch Disease Ontology (MONDO) [34] | 97,619 |
| Ontology for MIRNA Target (OMIT) [35] | 63,015 |
| Molecular Process Ontology (MOP) [36] | 57,355 |
| Name Reaction Ontology (RXNO) [37] | 57,330 |
| Human Phenotype Ontology (HP) [6] | 46,075 |
| Mammalian Phenotype Ontology (MP) [7] | 43,806 |
| Cell Ontology (CL) [38] | 34,685 |
| Ontology of Biological Attributes (OBA) [39] | 26,523 |
| Ontology of Adverse Events (OAE) [40] | 20,566 |

**Table 3.**
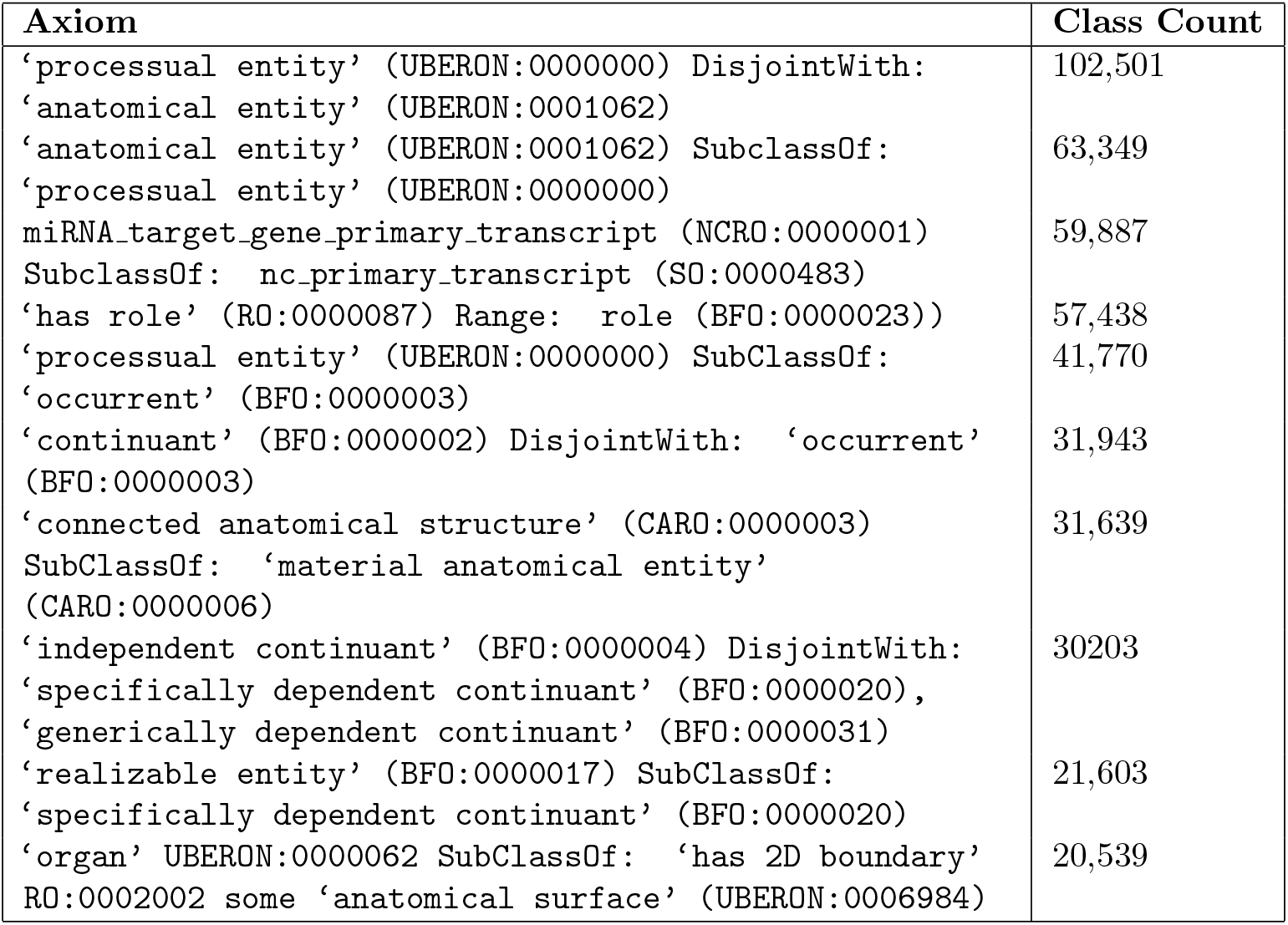
Top ten axioms accounting for the most hidden cases of unsatisfiability across OBO ontologies.

### Efficient ranking and repairing of axioms

Our algorithm for identifying the causes for unsatisfiability in ontologies builds upon a black-box algorithm for computing a justification for one unsatisfiable class. A justification is a minimal set of axioms which explain why the class is unsatisfiable. The black-box algorithm we employ creates an empty ontology containing only the class that is unsatisfiable; it then adds new axioms from the original ontology to it, until the class becomes unsatisfiable. Axioms that are not necessary for the class to become unsatisfiable are then removed using a backwards stepwise approach, eventually producing a minimal set of axioms that constitute a justification for the unsatisfiability of the class in the original ontology. Justification algorithms are usually used as debugging tools to direct ontology developers towards the causes of unsatisfiability. For this reason, they are often integrated into ontology development environments such as the Protégé software [41].

The naive algorithm, for finding a minimal set of justifications that can be removed to repair all cases of unsatisfiability, uses the black box algorithm to compute justifications for all unsatisfiable classes in the ontology, and then removes the axiom that appears most frequently in the set of all justifications. Subsequently, it then repeats this step until all cases of unsatisfiability are solved. This algorithm works well if only a small number of classes are unsatisfiable, and the ontology is relatively small. However, our experiments revealed a very large number of unsatisfiable classes, some of which are in very large ontologies. In the most prolific case, the Unified Phenotype Ontology (UPHENO) contains 106,126 unsatisfiable classes, out of 133,480 classes total in the ontology. Such a large number of unsatisfiable classes makes the naive algorithm intractable. In the worst case, our black-box algorithm has to add all axioms from the ontology, and then remove all but one of these axioms in order to find a single justification for one class, leading to a time complexity of (*n · m*) where *n* is the number of axioms and *m* the number of unsatisfiable classes; since each step further involves computing satisfiability, which has cubic complexity in the number of classes (and relations) [17], it is obvious that the algorithm will not scale to large numbers of unsatisfiable classes.

We develop an improved algorithm for finding a small set of axioms to remove from an ontology to repair all cases of unsatisfiability by a consideration of the problem according to the hitting set problem.

In the theory of system diagnosis, we consider a series of conflict sets, each describing a conflicting set of system components – a subset of elements from a universal set of system components. A hitting set is one which intersects every conflict set, and the hitting set problem is the problem of computing all the minimal hitting sets for the conflict sets [42].

The problem is useful in cases where repairing or removing all of the elements in a hitting set would repair a system. The hitting set problem is equivalent to the set cover problem [43], and both problems are known to be NP-complete through reduction to the boolean satisfiability problem [44].

Our problem can be reduced to the hitting set problem, because an unsatisfiability justification can be considered as a conflicting set of axioms which can be resolved by removing one of its members from the ontology. To completely remove all axioms causing unsatisfiable classes in an ontology, all justifications must be resolved.

A hitting set of axioms to remove from the ontology to repair all axioms, therefore, must have a non-empty intersection with every unsatisfiability justification. The problem of finding all justifications for a single entailment in an ontology has previously been reduced to the hitting set problem, and then solved using Reiter’s Hitting Set Tree (HST) algorithm [45]. The problem we need to solve is similar, however we need to identify a hitting set of axioms that resolve *all* cases of unsatisfiability in the ontology instead of just the axioms that cause unsatisfiability of a single class.

We develop an algorithm that exploits the fact that classes transitively inherit unsatisfiability through subclass axioms; if *C* is unsatisfiable and the ontology contains *D* ⊑ *C* as an axiom, then *D* will also be unsatisfiable. Consequently, we prioritise resolving unsatisfiabilities for classes that have the largest number of (asserted) subclasses in the ontology; when we resolve the cause of such a class becoming unsatisfiable, we also resolve the inherited causes of unsatisfiability for their subclasses without explicitly needing to generate a justification for them. In the worst case, this optimisation step will have no effect, because any class may have multiple causes of unsatisfiability independent from its parent class. If that is the case, the performance would be equivalent to the naive algorithm described above. However, commonly, if we assume that there are only a small number of overall causes of unsatisfiability in the ontology, we will reduce the number of justifications generated significantly.

Our algorithm is shown in Figure 2. The algorithm takes an ontology *O* as input and determines the set of unsatisfiable classes in *O*, *υ*(*O*); the algorithm then removes from *υ*(*O*) all classes that have an asserted superclass in *υ*(*O*). This step ensures that for each cluster of unsatisfiability, the most general class within the ontology taxonomy is examined first. The algorithm then selects the group of classes with the highest number of directly asserted subclasses, and either generates justifications for all of these classes or for a random sample of them if the number of direct subclasses is above a threshold *n* (we select *n* = 25). The most frequently occurring axiom in these justifications is then removed, and the ontology is reclassified, to produce another set of unsatisfiable classes, upon which the process is repeated; the algorithm terminates when all unsatisfiable classes have been resolved.

**Figure 2.**
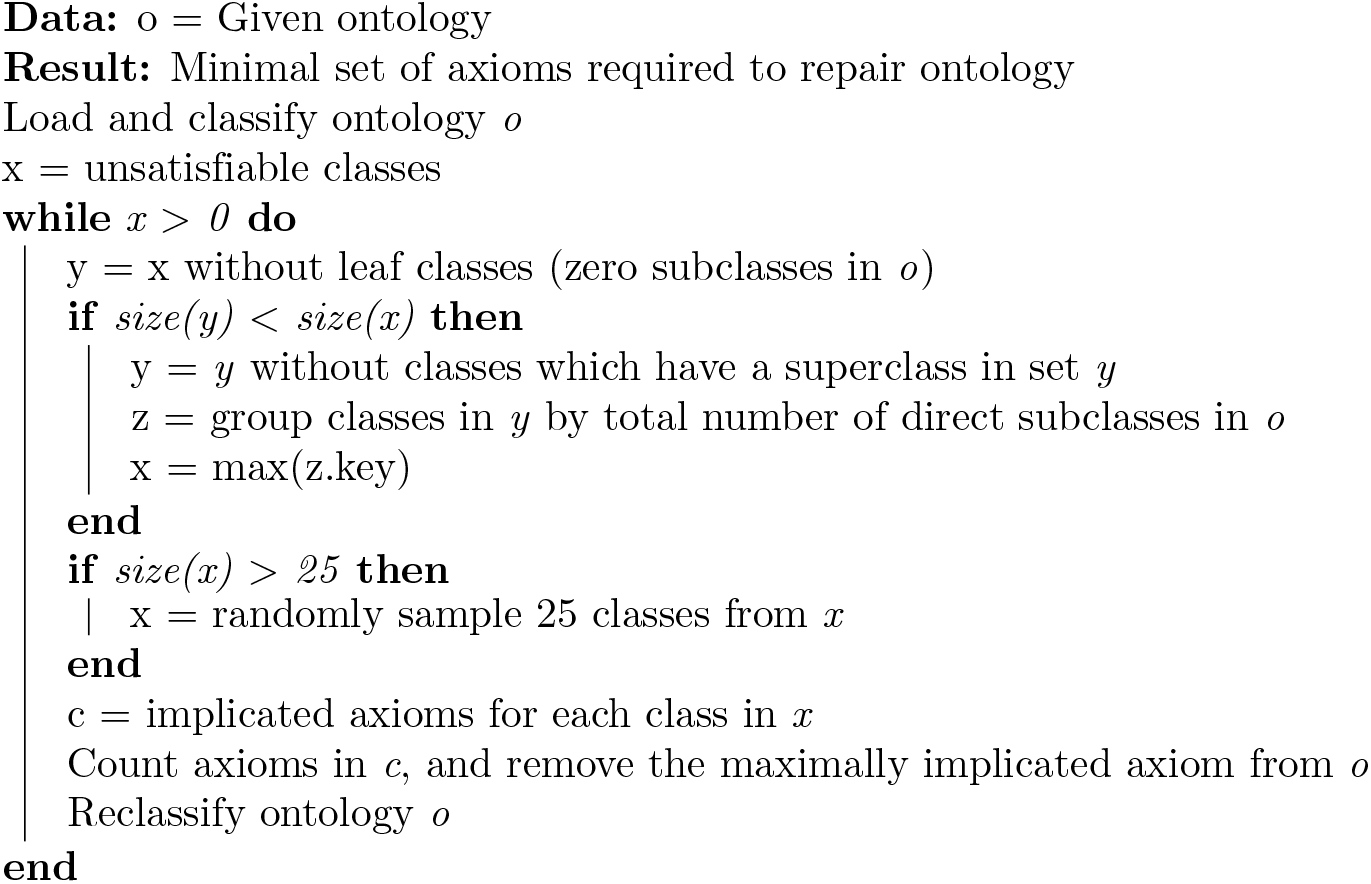
Algorithm for automatic diagnosis and repair of unsatisfiable classes in an ontology.

In the selection step, our algorithm uses asserted subclasses instead of inferred subclasses because each unsatisfiable class is an inferred subclass of all classes. It is possible that a class has more direct subclasses than another yet a fewer number of total subclasses; however, this effect is controlled by removing any classes with a superclass in the set of unsatisfiable classes *υ*(*O*).

Throughout execution of the algorithm, we record statistics on the set of classes that become satisfiable after the removal of each axiom. These statistics enable ontology developers to identify problematic axioms that affect groups of ontologies, and manually resolve them.

### Application to OBO Foundry

We applied our algorithm first to the merged OBO Foundry ontology, finding that two axioms could be removed to solve all cases of unsatisfiability:

1. `‘realizable entity’ (BFO:0000017) SubClassOf: ‘specifically dependent continuant’ (BFO:0000020)` with 599 classes repaired, and
2. `‘molecular entity’ (CHEBI:23367) SubClassOf: ‘material entity’ (BFO:0000040) with 37 classes repaired.`

These two axioms are members of the smallest set of axioms that suffices to remove all unsatisfiabilities. We could also consider the unsatisfiable classes as a result of violating disjointness axioms; in particular, all the unsatisfiable classes are also subclasses of two or more classes that are asserted to be disjoint. The removal of each of the subclass axioms above solves multiple disjointness axiom violations. For the first axiom that contributes to the most unsatisfiable classes, the classes it accounts for each violate one or more of these three different disjointness axioms:

1. `‘independent continuant’ (BFO:0000004) DisjointWith: ‘specifically dependent continuant’ (BFO:0000020)`
2. `DisjointClasses: ‘independent continuant’ (BFO:0000004), ‘specifically dependent continuant’ (BFO:0000020), ‘generically dependent continuant’ (BFO:0000031)`
3. `‘continuant’ (BFO:0000002) DisjointWith: ‘occurrent’ (BFO:0000003)`

The second case is affected by two disjointness axioms:

1. `‘independent continuant’ (BFO:0000004) DisjointWith: ‘specifically dependent continuant’ (BFO:0000020)`
2. `DisjointClasses: ‘independent continuant’ (BFO:0000004), ‘specifically dependent continuant’ (BFO:0000020), ‘generically dependent continuant’ (BFO:0000031)`

The two disjointness axioms shown for the second case are included in the three axioms shown for the first set, and the disjointness axiom between `independent continuant` and `specifically dependent continuant` is a consequence of the others. In total, therefore, three disjointness axioms account for all cases of hidden unsatisfiability throughout the OBO ontologies. Removing the subclass axioms removes fewer axioms and solves the cases of unsatisfiability because they prevent classes from violating multiple disjointness axioms. For example, in the case of removing the subclass relationship between `molecular entity` (`CHEBI:22367`) and `material entity` (`BFO:0000040`), some subclasses of `‘molecular entity’` violate the first disjointness axiom and some violate the second. By removing the subclass axiom, however, molecular entities are no longer subclasses of the parent class of `material entity, independent continuant` (`BFO:0000004`), for which two disjointness axioms are asserted.

Among the wider set of OBO ontologies we found that a set of only 117 axioms could be removed from ontologies to solve all unsatisfiability for all 866,494 unsatisfiable classes. Of these, 51 involved a BFO class. Figure 3 shows the top ten axioms ranked by the number of unique unsatisfiable classes they are responsible for repairing when removed, while the full set of axioms is available in the Github repository associated with this experiment.

**Figure 3.**
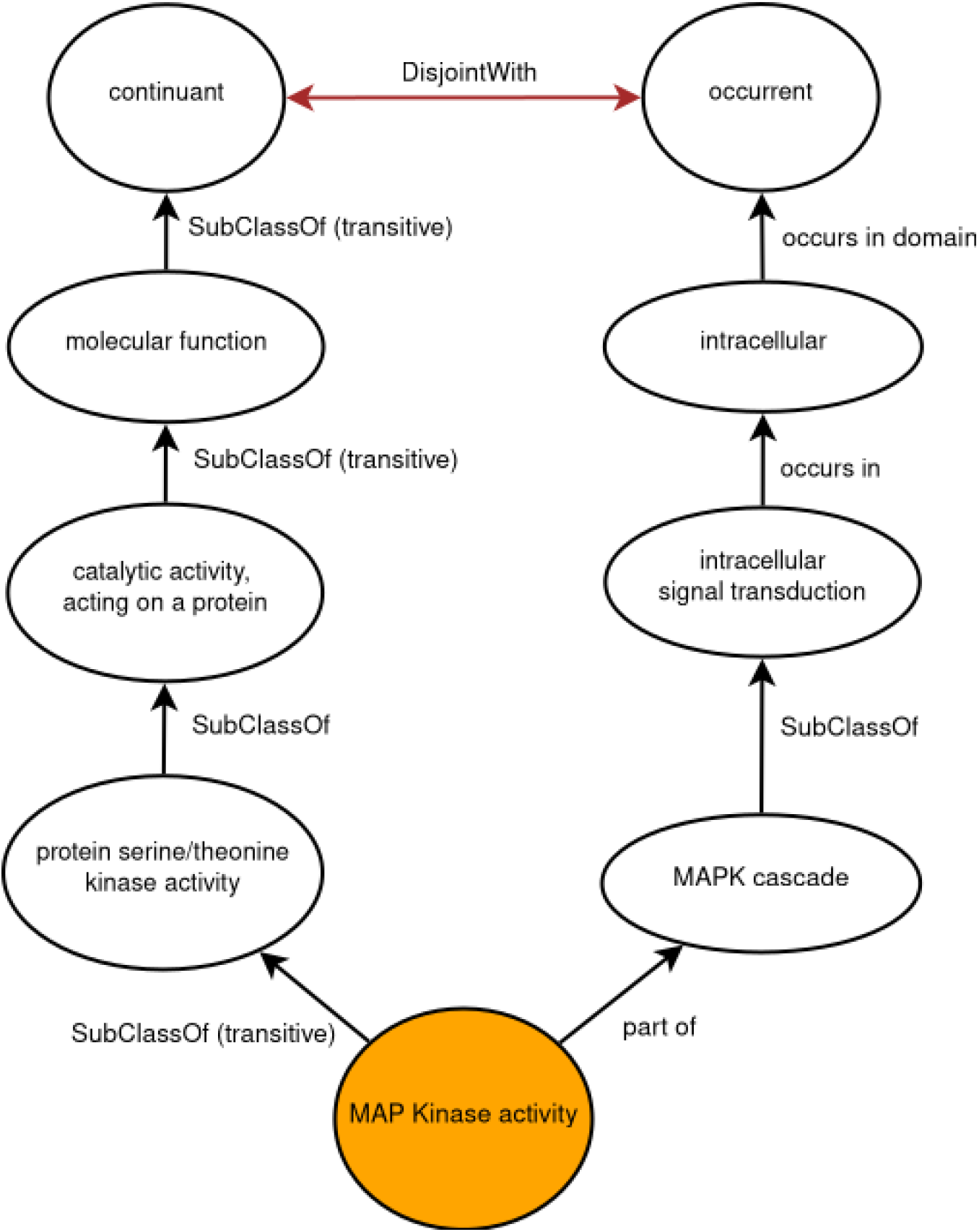
MAP Kinase unsatisfiability in the OBO Foundry meta-ontology represented as a graph.

### Inconsistency Analysis

Our experiments identify contradictions that lead to unsatisfiable classes in the OBO ontologies and highlight the axioms that can be removed to solve most cases of unsatisfiability. Our experiments further reveal which disjointness axioms are most frequently violated. However, merely removing the axioms does not necessarily resolve the underlying issues in how domain knowledge is modeled.

For example, although 599 unsatisfiable classes are repaired in OBO Foundry ontologies by removing the subclass axiom, `‘realizable entity’` (`BFO:0000017`) `SubClassOf: ‘specifically dependent continuant’ (BFO:0000020)`, this does not entail that this axiom, or the disjointness axioms it is related to, are themselves incorrect. Instead, the unsatisfiable classes arise through the different, mutually exclusive, uses of these classes by more specific axioms. In particular, 87 of these 599 classes are `MAP kinase activity` (`GO:0004707`) and its subclasses. The violated disjointness axiom is the fundamental BFO distinction between `continuant` (BFO:0000002) and `occurrent` (`BFO:0000003`). A continuant is something that is present as a whole at a time point and maintains its identity over time while an occurrent unfolds through time and has temporal parts [46]. They are often used in biomedical ontologies to refer to material entities and processes, respectively.

As shown in Figure 3, `MAP kinase activity` is a subclass of `continuant` (indirectly through several other classes) by means of being a `molecular function`. It is also a subclass of `part of` some `MAPK cascade`, which is a subclass of `intracellular signal transduction`. This class stands in an `occurs in` relationship with `intracellular`. The object property `occurs in` contains a restriction of its domain, asserting that something that `occurs in` something else must be an `occurrent`. Consequently, `MAPK cascade`, a kind of `intracellular signal transduction`, is an `occurrent`.

Then, because `MAP kinase activity is part of` a `MAPK cascade`, it too is an `occurrent`. The reason for this is that the `part of` (`BFO:0000050`) relationship must be between two things of the same kind; its description states “two distinct things cannot be part of each other” which is enforced by assertions in RO that state `occurrent` is a subclass of `part of` only `occurrent`, and continuant is a subclass of `part of` only `continuant`. This means that the reasoner infers from the assertion that `MAP kinase activity` is a part of `MAPK cascade`, that it too must be an `occurrent`. Therefore, `MAP kinase activity` must be both a `continuant` and an `occurrent`, which is the source of its unsatisfiability.

In addition to the 87 classes that due to the axioms related to `MAP kinase activity`, all 599 unsatisfiable classes that can be removed by removing the `‘realizable entity’` (`BFO:0000017`) `SubClassOf ‘specifically dependent continuant’` (`BFO:0000020`) axiom are subclasses of the class description:

- `‘molecular function’ and ‘occurs in’ some ‘intracellular’`

This is fundamentally the same cause of unsatisfiability as `MAP kinase activity`: that is they are subclasses of `continuant` via `molecular function`, and `occurrent` via being something or a part of something that `occurs in intracellular`. There are actually 1,306 total classes which are subclasses of `‘occurs in’ some ‘intracellular’`, but 707 of these are not subclasses of continuant and are therefore not unsatisfiable.

These contradictions are not revealed by an automated reasoner used with the Gene Ontology alone, because the Gene Ontology imports `occurs in` (`BFO:0000066`) from the Relation Ontology using MIREOT, without the axioms of the Relation Ontology. Consequently, the axiom that asserts the domain of `occurs in` is not imported. The contradiction is revealed when the ontologies are combined and the imported class is extended with the restrictions declared in its original definition.

The long chain of inferences required to detect this unsatisfiability explains why it is easy for an ontology developer to assert a contradictory axiom, especially when the full set of axioms is not available to a reasoner during ontology development. The shared inheritance of continuant and occurrent are hidden behind several layers of subclass axioms and domain and range restrictions on object properties. Furthermore, colloquially, there may also be occasional confusion between parthood and participation in a process [47]. The problems could be fixed without any removal of axioms by using the `participates in` (`RO:0000056`) or `has participant` (`RO:0000057`) relations instead of the `part of` relations in some axioms [48].

Indeed, many of the axioms that were highlighted for removal imply issues deriving from improper use of BFO. For example, in the OBO ontologies experiment, 57,438 classes were made satisfiable by removing the restriction that the role a class has must be a kind of role.

All tools described in this paper, including those to obtain, merge, analyse, and repair ontologies, as well as the full results of the experiment, and tools to recreate the experiment, are available at https://github.com/bio-ontology-research-group/UNMIREOT.

## Discussion

We have identified a high prevalence of hidden unsatisfiability throughout a major biomedical ontology ecosystem, the OBO ontologies. These ontologies include widely used ontologies that form a crucial part of the bioinformatics infrastructure. We also developed a novel algorithm that can diagnose incoherent ontologies by identifying a small set of axioms that resolve all cases of unsatisfiability. We demonstrated this across the OBO Foundry, and found that relatively few axioms can be removed to resolve all unsatisfiable classes. Nevertheless, the fact that many of the axioms removed belong to BFO, the upper-level ontology that most OBO ontologies use as a foundation, indicates that this ontology is not used consistently throughout all ontologies. Also of note is that several ontologies were inconsistent when combined with the set of OBO Foundry ontologies. These ontologies likely had similar problems to the other ontologies we examined, but actually included instances of the unsatisfiable classes – turning an incoherent ontology into an inconsistent one. Our algorithm reveals that it suffices to remove or change 117 axioms to repair all issues we identified; while our algorithm can automatically remove these axioms, the number of problematic axioms is small enough for them to be manually investigated; this sets out a way forward towards a logically consistent set of biomedical ontologies.

While our algorithm removes a minimal set of axioms to make an ontology coherent, it does not repair the root cause of the contradiction. In one case we showed that a large number of unsatisfiable classes in the Gene Ontology were caused by a mistaken use of a parthood relationship. This cause for unsatisfiability was complex, but would have been revealed by an automated reasoner had the axioms of MIREOT-ed classes been included. This indicates that the unconstrained use of MIREOT has introduced a new challenge for ontology interoperability, which must now be addressed. The question remains, however, of how best to balance the challenges of developing ontologies with the hardware resources and tools available, while at the same time maintaining consistency and interoperability between ontologies. Our results illustrate how the unMIREOT tool can be used to help ontology developers identify problematic axioms in their ontologies, and explore them to diagnose causes of contradiction.

While we have shown that there are large clusters of unsatisfiability across the OBO Foundry, it is unclear whether or to what extent these issues are affecting ontology-based analysis techniques. Incorrect inferences could affect the results of gene enrichment analysis, inter-ontology phenotype mapping, semantic similarity tasks, or any analysis that relies on ontology axiomatisation. In the future, we intend to explore this by implementing a reference task that relies on multiple combined ontologies, and comparing the performance before and after repairing the unsatisfiable classes.

While ontologies can be repaired by the unMIREOT approach, and examination of its output can help to identify the root cause of unsatisfiability, this can still be a time consuming and complicated process. It is possible that algorithmic tools could be developed to aid ontology developers in identifying the most informative cause of the inconsistency, or instead to create a set of minimally destructive axioms to remove from the ontologies.

One approach to preventing contradictions from entering ontology releases in the future is the the use of full ontology inclusion and testing during the development process, as part of an integration testing process. It would be possible to incorporate the unMIREOT tool in such a workflow or ontology release tool [49]. The OBO ontologies use a shared central build system which can be configured to validate ontologies against scripts that check for problems. By using a powerful build server to combine ontologies with the ontologies they refer to and check for inconsistencies before release, developers would be able to continue to use MIREOT while ensuring continuing compatibility.

It is also possible that either the MIREOT or OBO guidelines should be revised, to include more information in a class reference. Including more axioms related to referenced classes would allow for local consistency checking with an automated reasoner. Because many axioms are inherited, and restrictions are placed transitively, the axioms of an entire ontology or at least a derived module would need to be imported. This could be recommended in the case of small, high-level ontologies such as BFO and RO, which should not cause performance or space issues. Without actually including the ontology in the imports closure, however, it would not solve the problem of sourcing ontologies becoming out of date with the ontologies they reference.

## Author’s contributions

RH and LTS conceived of the study and experimental design. LTS performed all experiments and implemented the software. LTS drafted the manuscript. RH and GVG supervised the project. All authors revised and approved the manuscript for submission.

## Acknowledgements

GVG and LTS acknowledge support from support from the NIHR Birmingham ECMC, NIHR Birmingham SRMRC, Nanocommons H2020-EU (731032) and the NIHR Birmingham Biomedical Research Centre and the MRC HDR UK (HDRUK/CFC/01), an initiative funded by UK Research and Innovation, Department of Health and Social Care (England) and the devolved administrations, and leading medical research charities. The views expressed in this publication are those of the authors and not necessarily those of the NHS, the National Institute for Health Research, the Medical Research Council or the Department of Health.

RH and GVG were supported by funding from King Abdullah University of Science and Technology (KAUST) Office of Sponsored Research (OSR) under Award No. URF/1/3790-01-01.

